# Genome-wide *cis*-decoding for expression designing in tomato using cistrome data and explainable deep learning

**DOI:** 10.1101/2021.06.01.446518

**Authors:** Takashi Akagi, Kanae Masuda, Eriko Kuwada, Kouki Takeshita, Taiji Kawakatsu, Tohru Ariizumi, Yasutaka Kubo, Koichiro Ushijima, Seiichi Uchida

**Affiliations:** Graduate School of Environmental and Life Science, Okayama University, Okayama 700-8530, Japan; JST, PRESTO, Kawaguchi-shi, Saitama 332-0012, Japan; Department of Advanced Information Technology, Kyushu University, Fukuoka 819-0395, Japan; Division of Biotechnology, Institute of Agrobiological Sciences, National Agriculture and Food Research Organization, Tsukuba, Ibaraki 305-8602; Faculty of Life and Environmental Sciences, University of Tsukuba, Gene Research Center, Tsukuba, Japan

## Abstract

In the evolutionary paths of plants, variations of the *cis*-regulatory elements (CREs) resulting in expression diversification have played a central role in driving the establishment of lineage-specific traits. However, it is difficult to predict expression behaviors from the CRE patterns to properly harness them, mainly because the biological processes are complex. In this study, we used cistrome datasets and explainable convolutional neural network (CNN) frameworks to predict genome-wide expression patterns in tomato fruits from the DNA sequences in gene regulatory regions. By fixing the effects of *trans-*elements using single cell-type spatiotemporal transcriptome data for the response variables, we developed a prediction model of a key expression pattern for the initiation of tomato fruit ripening. Feature visualization of the CNNs identified nucleotide residues critical to the objective expression pattern in each gene and their effects, were validated experimentally in ripening tomato fruits. This *cis*-decoding framework will not only contribute to understanding the regulatory networks derived from CREs and transcription factor interactions, but also provide a flexible way of designing alleles with optimized expression.

## MAIN TEXT

*Cis*-regulatory elements (CREs) are noncoding short DNA sequences that are recognized by transcription factors (TFs, or *trans-*element). CREs play a central role in the regulation of gene expression. In the diversification of plants, including whole-genome duplication events, the evolution of CREs has made more rapid and substantial contributions to lineage-specific acquisition of representative traits than that of *trans-*elements^1-3^. This role of CREs also has been reported in animals^4-6^. Variation of gene regulatory regions or CREs also have had major impacts on the evolution of crops^7-9^, and a next-generation breeding platform with *cis*-editing has been proposed to allow expression fine-tuning^10-11^. However, unlike *trans-*elements, which have been studied extensively, little is known about the functions of CREs to properly harness them. This is mainly because of the structural complexity of the biological processes involved. A TF can bind to multiple and plastic motifs (or CREs)^12-13^, which makes it difficult to define the uniform motif sequences. Furthermore, even if the effects of *trans-*elements can be fixed, the gene expression pattern will be determined by a flexible combination of multiple CREs^14^, depending on their positional relationships.

Deep learning (DL) techniques that utilize convolutional neural networks (CNNs) have contributed to breakthroughs mainly in image diagnosis and natural language processing^15^. Unlike conventional machine learning, DL algorithms can automatically find flexible and complicated features. Although DL predictions had been a black box and difficult to explain, methods for feature visualization of DL predictions (often referred to as explainable AI) have been recently developed^16-17^. These methods have allowed the biological interpretation of DL predictions, thereby accelerating the application of DL techniques in plant biology^18-19^. DL methods also have been used to predict transcript regulatory regions^20-22^ and epigenetic marks, such as DNA methylation levels^23^, in genomic sequences. Importantly, the combination of explainable DL predications and high-throughput enrichment of TF-bound DNAs (e.g., by ChIP sequencing) has successfully produced high-quality predictions of CREs and enabled the identification of the nucleotide sequence motifs responsible for TF binding^24^. These findings suggested that, with a trained explainable DL model, DNA sequences could be encoded into CREs for each TF, and CREs could be decoded into the residues responsible for binding.

Cistrome databases constructed using protein binding microarray, and ChIP or DAP sequencing data have comprehensively accumulated short sequences that contain CREs. These databases cover most TF families in eukaryotes^12^, including Arabidopsis^13^ and other plant species^25^. The affinities of TF DNA binding domains nested into the same TF family are highly conserved across species^12,25^, which enabled interspecific annotation of the CREs in some CRE databases^25-26^. On the basis of these findings, we aimed to develop a DL framework to predict gene expression patterns from their promoter sequences (Fig. 1), (i) by predicting CREs in new promoter sequences using the large cistrome datasets from model plants, (ii) by constructing models to predict expression patterns from CRE arrays, and (iii) by identifying the key nucleotide residues responsible to the predicted expression patterns. We exemplified differential expressions in ripening tomato fruit to fine-tuning the gene expression patterns related to maturation/softening patterns, which has been a key issue since the 1980s^27-30^.

**Figure 1.**
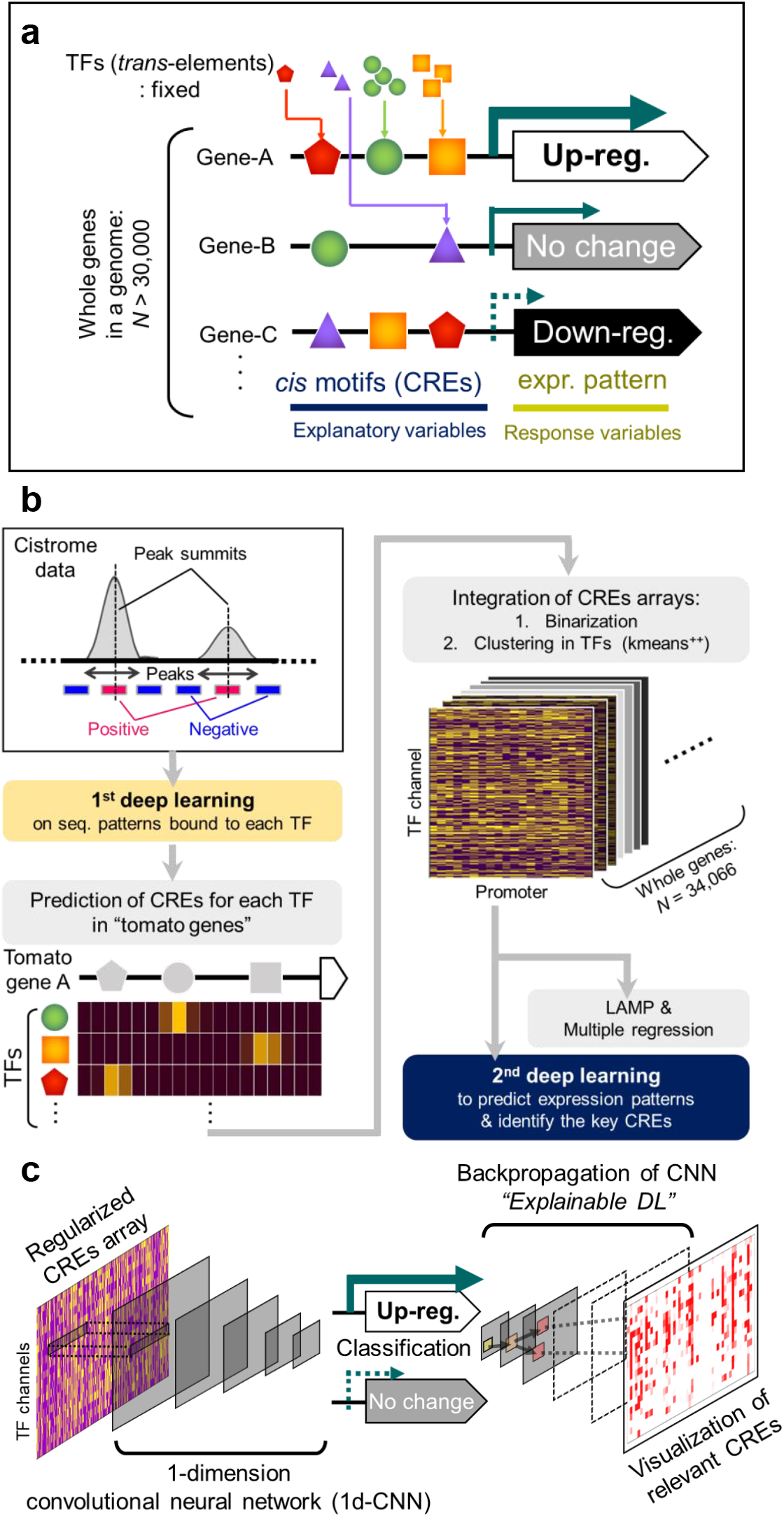
Prediction of gene expression patterns in a genome from *cis*-regulatory elements (CREs). **a**. Schematic model for the prediction of the expression patterns among all the genes in a genome. In a homogeneous cell line, the effects from *trans-*elements can be fixed among the genes. Then, expression patterns can be explained from flexible combinations of CREs (and potential epigenetic marks). **b**. Construction of the prediction model with two-step deep learning (DL) frameworks. Large Arabidopsis cistrome datasets^13^, which provide genome-wide transcription factor (TF)-binding peaks, were used in the first step (1^st^ deep learning) to predict CRE patterns for each TF. The resultant model was applied to the tomato genome sequences to predict CREs in the promoters of all the genes, to derive CRE arrays. For each gene, the CRE array was annotated with an expression pattern that was applied to the second step (2^nd^ deep learning) and also used for the multiple regression and LAMP analyses^14^. **c**. In the 2^nd^ deep learning step, the CRE arrays were trained with a one-dimensional convolutional neural network (1d-CNN) with the clustered TF channels to make a binary classification. With backpropagation of the CNN (explainable DL), the CREs or other nucleotide residues relevant to the objective expression class were visualized.

From the Arabidopsis DAP-seq (cistrome) dataset for 529 TFs, which covers most plant TF families^13^, bilateral 15-bp nucleotide sequences were extracted from the TF-binding peak summit as the positive tiles, and same length (31-bp) nucleotide sequences were randomly extracted out of the peak area as the negative tiles (Fig. 1b). We used only the high-confidence DAP-seq data with fraction of reads in peaks >0.05^13^, covering 370 TFs (Supplementary Table S1). The 31-bp sequences were converted into a one-hot array with four A/T/G/C channels^31^. There have been many powerful tools for motif discovery, such as MEME^32^ or deep learning (DL)-based techniques^24^, while we adopted a simple in-house fully connected DL model (see Methods for details), to rapidly find the residues relevant to the prediction by feature visualizations, and to directly connect to the following in-house 2^nd^ DL model which predicts expression behaviors, as discussed later. Most of the 370 TFs datasets had high classification abilities as indicated by the ROC (Receiver Operating Characteristic) curves (Fig. 2a, area under the curve (AUC) values = 0.956 ± 0.0022 on average, Supplementary Table S2). Their classification abilities were mostly comparable to those with the popular MEME motif discovery tool^32^, in recall, precision, and F1 scores on the same train/test sample sets (Fig. 2b). These methods showed high correlations in their classification abilities among the TFs, suggesting their performances depended substantially on the characteristics of the TFs and/or the quality of the cistrome data (Supplementary Fig. S1, Table S3). Two distinct feature visualization methods, Guided Grad-CAM and Layer-wise relevance propagation (LRP), consistently detected not only representative motifs as relevant residues, which have been well characterized in previous studies and registered in the databases^13^, but also motif variants that showed significant peaks in the DAP-seq, which were quite similar to the representative ones but contained minor substitutions or gaps (Fig. 2c, representing ABF2 as a bZIP TF to bind to G-box: (C)ACGT(G), Supplementary Fig. S2 for three other TF families). An advantage of using DL models is their flexibility in accepting these small variations. We applied the 370 trained DL models to the 1-kb promoter regions of all the genes in the tomato genome (ITAG 4.0^2^, N = 34,066 with qualified promoter sequences) to predict the CREs for each TF. The predicted CRE transitions were converted into binary arrays with a 0.8 confidence threshold per 10–50-bp bin, and used to cluster the TFs with K-means^++^ (Supplementary Table S4), to avoid the multicollinearity problems in the assessments. With *K*=50, which is a hypothetically optimal cluster number (Supplementary Fig. S3), 42 clusters contained mostly a single TF family (these clusters were named with the predominant TFs); the other eight clusters contained a variety of different TFs (Fig. 2d, Supplementary Table S5). For the subsequent analyses, the CRE arrays of the 50 TFs that were closest to the center pattern in each cluster, were used as representatives for the 50 clusters.

**Figure 2.**
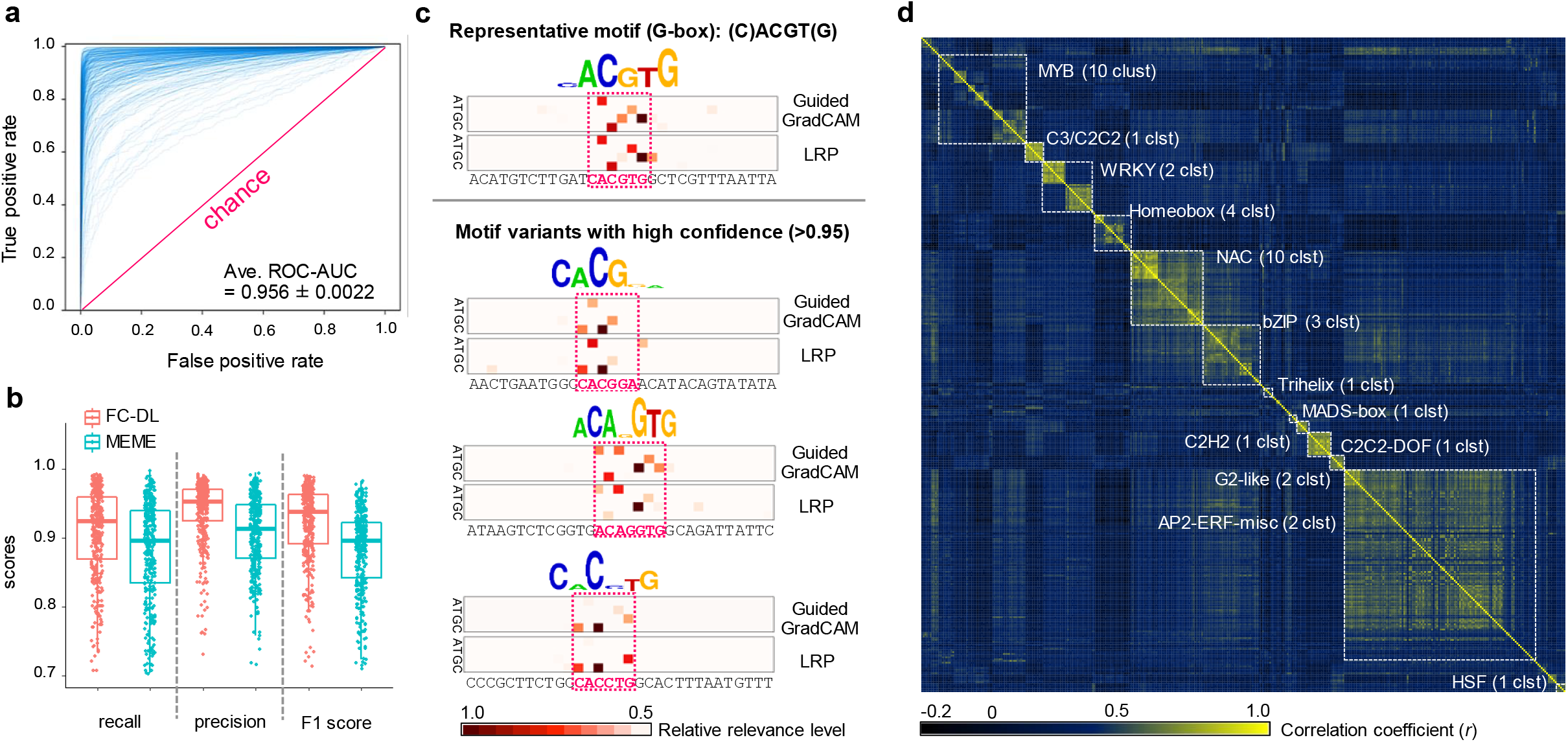
High-confidence prediction of variable *cis*-regulatory elements (CREs) and key nucleotide residues by deep learning (DL) **a**. Receiver Operating Characteristic (ROC) curves for the binary classification of transcription factor (TF)-binding and control sequences, for 370 TFs. The area under the curve (AUC) values ranged from 0.708 to 0.998 (average of 0.956). **b**. Prediction performance of the fully-connected deep learning model (FC-DL) and MEME (as used in O’Malley et al. 2016^13^). **c**. Nucleotide residues relevant to the prediction of CREs by the DL model, determined using two distinct feature visualization methods, Guided GradCAM and LRP. Relevance levels in the putative CREs (in the pink dotted squares) are reflected in the height of the nucleotide logos. *ABF2*-binding sequence tiles with high-confidence (>0.95) for the prediction are represented. The prediction model properly highlighted the residues consistent with the physiologically validated representative motif (C)ACGT(G), which is a bZIP-binding G-box core motif^46^. Furthermore, the same model detected motif variants, including minor gaps or substitutions. **d**. Correlation matrix for the CREs of the 370 TFs, with clustering by K-means^++^ (K=50). Each cluster was constituted mostly of TFs from the same family (see Supplementary Table S5 for detail).

We used a high-resolution spatiotemporal expression map of tomato fruit^33^ and focused on the gene expression patterns only in the pericarp, from “mature green (MG)” to “breaker (BR)” stages, which is a key transition for ripening initiation. In a transcriptome with heterogeneous cell lines, such as flower or leaf, the output is a mixture of multiple expression patterns derived from the expression of heterogeneous *trans-*elements (Supplementary Fig. S4). Instead, the extracted transcriptomes derived from a single (or homogeneous) cell type can fix the effect from *trans-*elements, thereby facilitating the construction of a precise model between CREs and genomic expression patterns (Fig. 1a). We focused on genes that were significantly upregulated or downregulated from MG to BR (defined as BRup and BRdown, respectively) (Fig. 3a, FDR <0.1 with DESeq2 analysis, >1.7-fold changes, RPKM >1). Total 34,066 arrays for all the genes in the tomato genome with the 50 described TF channels (refer to Fig. 1b), were trained with in-house one-dimensional CNN models (see the Methods for the detailed setting) to classify the expression patterns into the binary categories. The models for classification of the BRup and BRdown achieved averaged ROC-AUC values of 0.702 and 0.636, respectively (Fig. 3b, with four-fold cross-validation, Supplementary Fig. S5 for ROC and learning curves). Notably, these would be still far from perfect prediction, while the gene expression patterns in tomato fruit are affected not only by DNA sequence-based CRE variables but also by many kinds of epigenetic marks or chromatin folding^34-36^, even when the *trans-*elemental effects are fixed. Furthermore, indirect TFs binding, which depends on interactions between TFs, would have substantial effects on expression pattern^13^. Thus, future implementation of a multiple-inputs model that can consider also the epigenetic variables and TFs interactions may improve the performance. In the classification model for BRup, with the higher prediction performance than BRdown, the confidence distributions of the positive (or upregulated in BR) and the negative (control) genes were statistically separated (Fig. 3c, P <2.2e−16), whereas they were not significantly correlated to the expression levels (RPKM) or biases between MG and BR (Supplementary Fig. S6). The positive genes with the highest 10% confidence (*N* = 297), were significantly enriched with gene ontology (GO) terms involved in ethylene signals compared with those of all the positive genes (Fig. 3d). In climacteric fruit crops, including tomato, ethylene signals are thought to be the key pathway for ripening, suggesting that this model would be competent for designing the expression profile to tune the ripening process.

**Figure 3.**
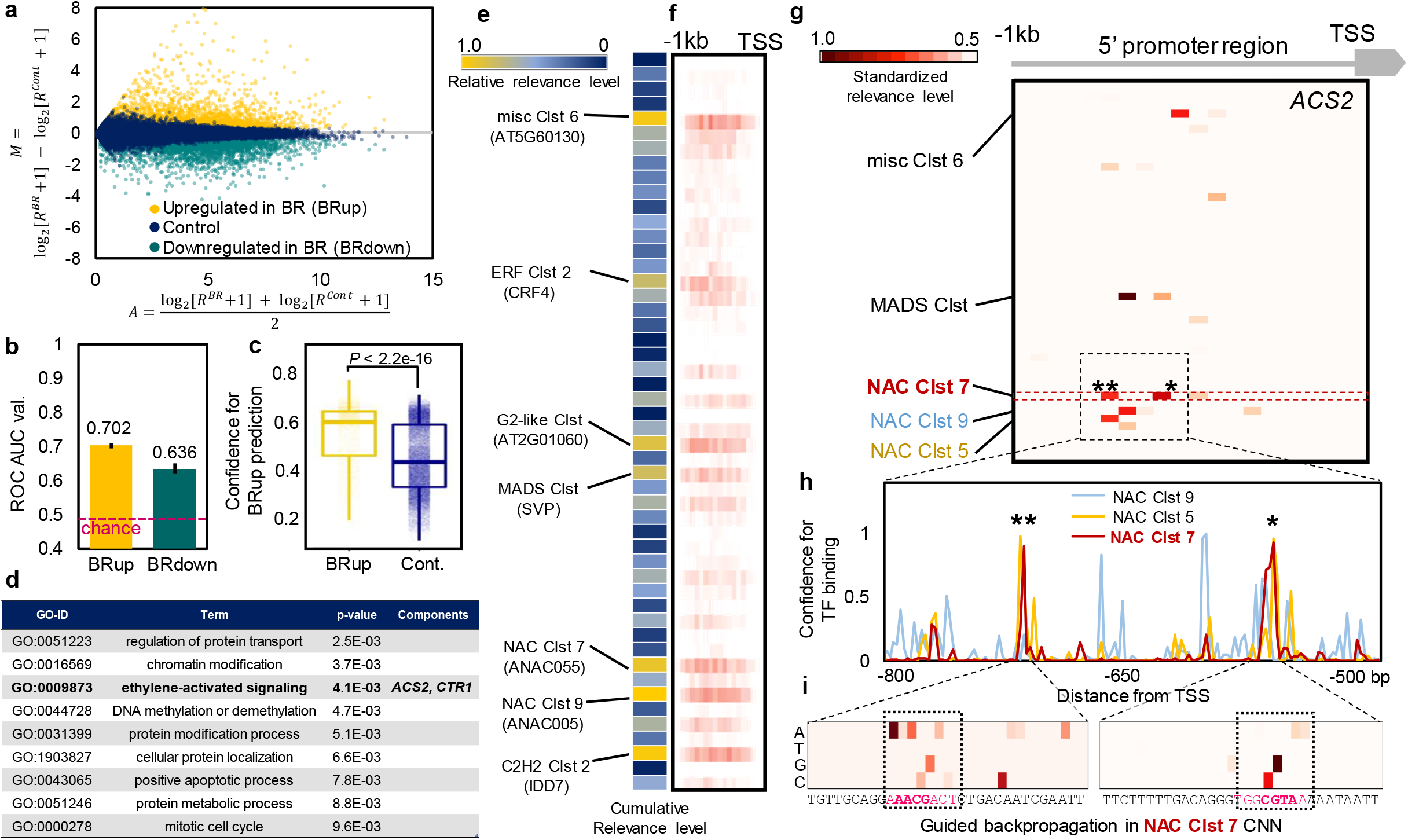
Prediction of the gene expression patterns critical to tomato fruit ripening initiation by deep learning, and visualization of their key cis-elements. **a**. MA plot for the genes expressed in the mature green (MG) and breaker (BR) stages of ripening tomato fruit. Genes significantly upregulated in BR (*N* = 2,967, defined as “BRup”) and downregulated in BR (*N* =3,098, defined as “BRdown”) are shown in orange and deep green, respectively. **b**. Performances (ROC-AUC values) for binary classification of the BRup or BRdown against the control category. Averaged ROC-AUC values were calculated from four-fold cross-validations. Bars indicate standard errors (SE). **c**. Confidence distribution in BRup prediction. Actual BRup genes exhibited substantially higher confidences than in the control genes (*P* < 2.2e-16). **d**. Gene-ontology (GO) terms significantly enriched in the genes with the highest 10% confidence in the BRup category. **e**–**f**. Predicted cumulative relevance levels for the genes with the highest 10% confidence in the BRup category (**e**) and per position in 50-bp bins (**f**). Of the 50 channels recognized by each TF cluster, the seven with high relative relevance levels (>0.7) are highlighted. The center TF for each cluster is in parenthesis. **g**–**i**. Identification of the CREs responsible for the BRup in the promoter region of the *ACS2*. With guided backpropagation on the model for BRup prediction, five channels showed high relevance levels (**g**). NAC Clst 7, the channel with the highest cumulative relevance level, showed two major relevant bins that corresponded to the high-confidence TF-binding regions, as indicated by the asterisks (* and **) (**h**). Further guided backpropagation on the model for CRE prediction from the promoter sequences tiles (the 1^st^ deep learning step, see Fig. 1b), the nucleotide residues responsible for the two TF-binding regions were detected (**i**). The most relevant residues were localized on the hypothetical NAC-binding motifs indicated by dotted squares.

To identify CREs relevant to the prediction of upregulation in BR, we applied a feature visualization method, Guided backpropagation, to the 297 high-confidence genes. Cumulative relevance levels were enriched in the channels recognized by NAC, C2H2, MADS-box, G2-like, and ERF TF clusters (Fig. 3e-f). This result was supported by the conventional multiple regression test and LAMP analysis (http://a-terada.github.io/lamp/), which can detect combinations of CREs that contribute to differential expression^14^ based on CRE numbers (Supplementary Table S8 and S9), although the CRE positions were not considered in these methods. Importantly, these high-relevance 5 TF families included key genes for the initiation of tomato fruit ripening, such as *NON-RIPENING*^37^ (*NOR*, NAC family), *SlZFP2*^38^ (C2H2 family), *RIPENING-INHIBITOR*^29^ (*RIN*, MADS-box family), and some *ETHYLENE RESPONSE FACTORs* (*ERF*s)^39-41^. This suggested that our *in silico* feature prediction properly reflected the actual physiological relationships, and may be applicable to estimate *trans-*factors (or upstream regulatory networks) directly involved in the objective expression patterns. In exemplifying key ethylene-producing enzymatic genes, the aminocyclopropane-1-carboxylic acid synthase 2 gene (*ACS2*) from the ethylene signal-related genes with high confidence (Supplementary Table S10), relatively higher relevance was localized in the channels recognized by three TF clusters, NAC (NAC Clusters 5, 7, 9), MADS-box (MADS Cluster), and a miscellaneous (misc Cluster 6) (Fig. 3g). Tomato *NOR* TF, which potentially controls the upregulation of *ACS2* in BR fruit^42^, was phylogenetically nested into the NAC Cluster 7 (Supplementary Fig. S7). NAC Cluster 7 had two high-confidence binding peaks in *ACS2*, with positions that were consistent with the high relevance bins (Fig. 3h). Further feature visualization with Guided backpropagation in the first DL model, which predicted CREs from the DNA sequence tiles (see Fig. 1b), localized high relevance on a few nucleotide residues, consistent with the hypothetical NAC-binding sequences (Fig. 3i).

To experimentally validate the prediction of the DL models, we artificially mutated the nucleotide residues relevant to the upregulation in BR (p*ACS2mut*). The p*ACS2mut* promoter sequences showed a substantial reduction in confidence for the two NAC Cluster 7 binding peaks (Fig. 4a), resulting in low confidence for upregulation in BR (Fig. 4b, from 69% for the intact p*ACS2* to 17% for p*ACS2mut*). A transient reporter assays in tomato fruits in MG, BR, and further ripened light red (LR) stages, with the luciferase reporter under the control of the p*ACS2* or p*ACS2mut* (Fig. 4c for the constructs), showed that the p*ACS2mut* was significantly less upregulated than the intact p*ACS2* in a BR stage-specific manner (Fig. 4d, *P* =1.1e-5 for BR, 0.98 for MG and 0.64 for LR, Student’s *t*-test). This result suggested that the targeted (or mutated) nucleotide residues would be critical for upregulation of *ACS2* from MG to BR, which was consistent with the prediction of the DL model. To further test the activation ability of p*ACS2* and p*ACS2mut* by NAC Cluster 7 TF, transient reporter assay in *Nicotiana benthamiana* leaf was conducted with luciferase reporters under the control of the p*ACS2* or p*ACS2mut* and the effector of constitutively expressed tomato *NOR* (p35S-*NOR*, see Fig. 4d). The mutations in p*ACS2* abolished activation by *NOR* (Fig. 4f, *P* =4.0e-6). The consistent results were obtained also with GFP reporters in *N. benthamiana* (Supplementary Fig. S8), and in previous reports focusing on *NOR*- and *NOR*-like TFs functions in ripening tomato^42,43^. Together, all the wet experimental results were consistent with the predictions from the DL models.

**Figure 4.**
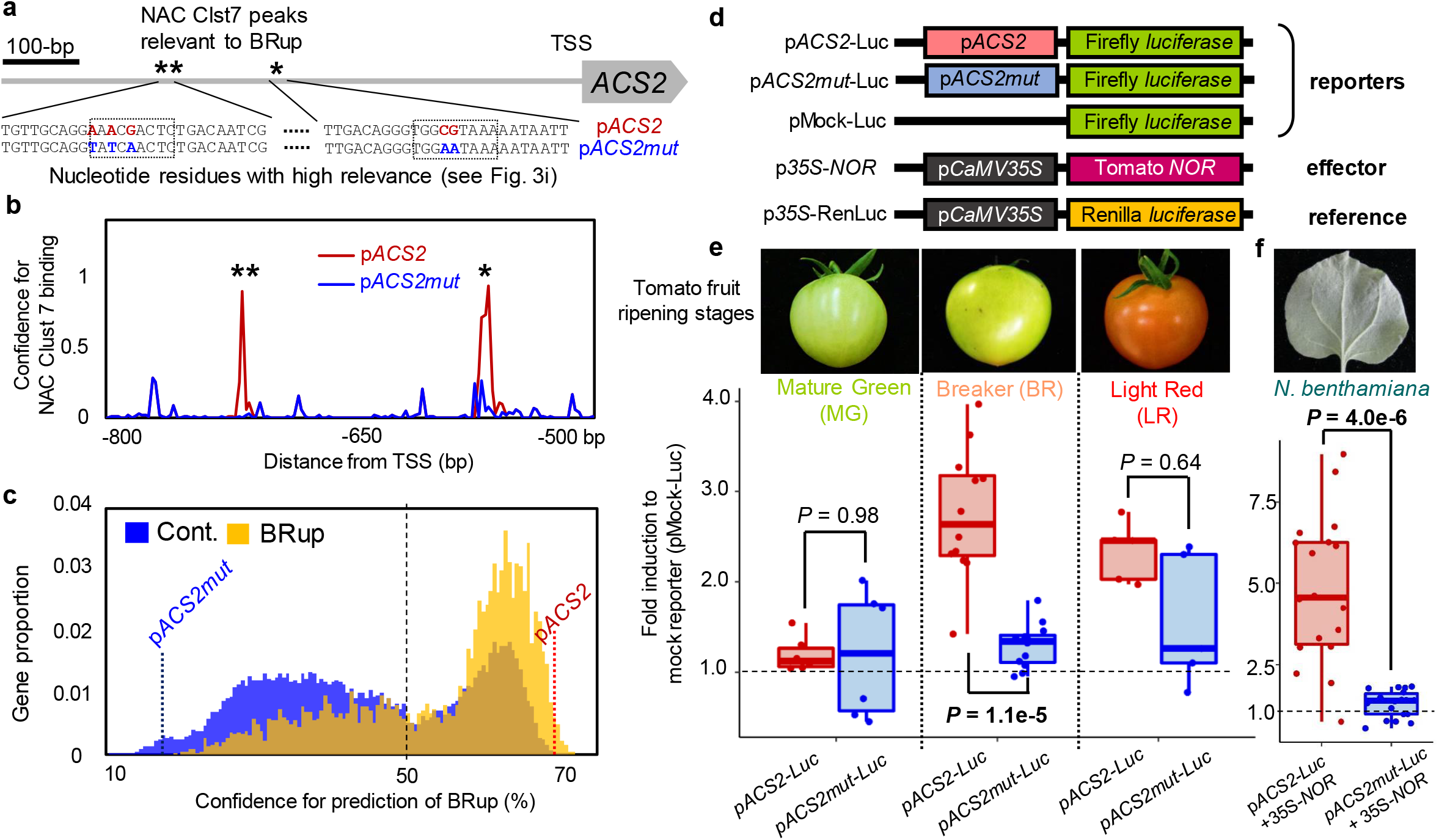
Experimental validations for the *cis*-decoding by deep learning. **a**. Point-mutations were artificially made on the residues with high relevance to the DL prediction (see Fig. 3i) in the 1-kb promoter of *ACS2* (pACS2, in red), deriving the mutated allele named p*ACS2mut* (in blue). **b-c**. p*ACS2mut* showed a substantial reduction in the confidence for the NAC Clst 7 binding prediction (**b**), and for the BRup prediction (Conf. = 69% for p*ACS2*, and 18% for p*ACS2mut*) (**c**). **d**. Constructs for transient reporter assays. **e**. Dual-luciferase transient reporter assay in ripening tomato fruits. In the mature green (MG) stage, p*ACS2* and p*ACS2mut* showed no significant differences (*P* = 0.98), and only slight activation compared with that of the mock reporter. In the breaker (BR) stage, p*ACS2* showed stronger activation than in the MG stage, whereas *ACS2mut* was substantially less activated (*P* =1.1e-5, Student’s *t*-test). In the light red (LR) stage, both p*ACS2* and p*ACS2mut* were activated in comparison to the mock, while they showed no statistical differences (*P* =0.64). **f**. Transient reporter assay with *Nicotiana benthamiana* for activation of p*ACS2* and p*ACS2mut* alleles by a tomato ripening key factor, *NOR*, nested in the NAC Clst 7. Constitutive expression of tomato *NOR* could induce p*ACS2* activation, while p*ACS2mut* was not substantially activated (*P* =4.0e-6, Student’s *t*-test).

In this study, *ACS2* was exemplified, while other representative genes involving fruit ripening initiation, such as *ACS4, Polygaracturonase* (PG), and *Pectin lyase* (PL), also exhibited high confidence for the BRup prediction (Table S10), in which visualized responsible CREs are consistent with previous studies (Supplementary Fig. 9). This *cis*-decoding framework will not only be applicable to characterization of the regulatory networks derived from CREs and transcription factor interactions, but also to designing alleles with optimized expression (Fig. 5). Once a good model for predicting expressions from the CRE array is constructed, feature visualization steps would find nucleotide residues responsible for objective expression. Artificial mutation or modification of the responsible residues would efficiently invent a new expression pattern, which can be predicted using the two-step DL models *in silico*. If an optimized expression is predicted, the flexible genome-editing system with CRISPR-Cas9^44^ can be used to design the allele for optic expression, as was partially shown in our modification of the *ACS2* promoter. In crops such as rice, tomato, grape, and apple, natural variations in the CREs have had major impacts on the development of the novel traits and phenotypic diversity that are critical for their qualities^7-9,45^ (summarized in Li et al. 2020^11^). Learned from their historical blueprints, multi-aspect *cis*-engineering, which unlocks the current breeding limitations and finely tunes the traits sensitive to the expression balances, has been proposed in some crops^11^ and has been attempted based on random mutations with the CRISPR-Cas9 system^10^. Our *cis*-decoding methods with explainable DLs will contribute to the further development of these possibilities and accelerate their implementation.

**Figure 5.**
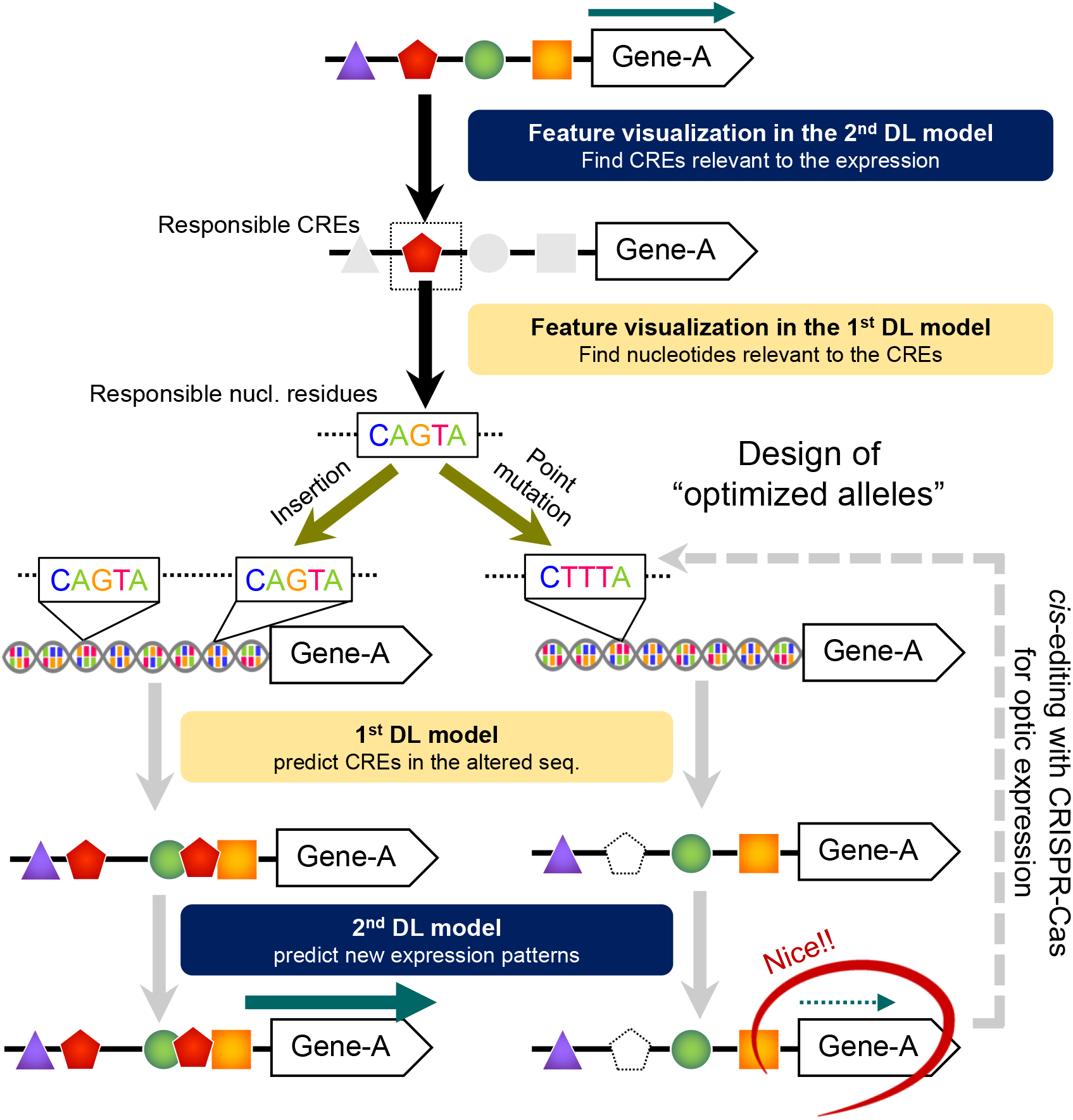
Model for expression design based on explainable deep learning (DL) If the objective expression patterns can be well predicted from *cis*-regulatory element (CRE) arrays, two-step feature visualization in the prediction models (or the 2^nd^ and then 1^st^ DL models, see Fig. 1b) will allow identification of the nucleotide-scale factor(s) responsible for the expression pattern. Randomization of the responsible residues can derive potentially unlimited variations for the objective expression pattern, which can be easily predicted using the 1^st^ and 2^nd^ DL models. Once a desirable expression pattern is predicted, *cis*-editing with the CRISPR-Cas system may realize the design of the optimized allele.

## METHODS

### Mining of cistrome datasets

We downloaded the transcription factor (TF)-binding peaks in narrowPeak format (fraction of reads in peak ≥5%) from the Arabidopsis DAP-seq datasets for 529 TFs^12^ (http://neomorph.salk.edu/dev/pages/shhuang/dap_web/pages/index.php). Bilateral 15-bp sequences from the peak summits (and their reverse complementary sequences) were extracted as the DNA tiles that included TF-binding sites (positive tiles for the DL classification). The 31-bp tiles out of the peak area were randomly extracted as the negative control tiles that included no TF-binding sites. The numbers of the positive/negative tiles applied to the DL classification are summarized in Supplementary Table S1.

### Mining of transcriptomic datasets of ripening tomato fruit

We downloaded mRNA-seq datasets of pericarp at the five typical ripening stages (mature green (MG), breaker (BR), pink, light red, and ripening red) in fastq format from the spatiotemporal expression map of tomato fruit^33^. The mRNA reads were mapped to the tomato reference protein-coding sequences (CDS) set (ITAG4.0, ftp://ftp.solgenomics.net/tomato_genome/annotation/ITAG4.0_release/) using BWA with the default settings. The mapped reads were counted to calculate the RPKM (reads per kilobase per million mapped reads). Differentially expressed genes between MG and BR were detected using DESeq2. Genes that were upregulated or downregulated from MG to BR (BRup and BRdown genes, respectively) with FDR <0.1 and RPKM >1.0 (Supplementary Table S6-7), were used for the DL classification analyses.

### Deep learning models for predicting the *cis*-regulatory elements (CREs) from cistrome datasets

For each TF, we randomly selected 20% of the positive and negative 31-bp tiles from the cistrome datasets for the test dataset. We allocated 70% and 30% of the remaining tiles to the training and validation datasets, respectively. These two datasets were used in a fully-connection model that had three layers (see “FC-DNA_H5-ROC_ConfMatrix.py” in https://github.com/Takeshiddd/Genome-wide-cis-decoding-for-expression-designing-in-tomato-with-cistrome-and-explainable-deep-lear) and was constructed with the sequential API model of Keras 2.2.4 (https://keras.io/). We set the class weight option (“class_weight” in Keras) with the bias in the sample numbers in the two classes. We uniformly set the epoch = 15, the learning rate = 0.001, and used the Adaptive Moment Estimation (Adam) optimizer among the 370 TFs datasets. The performances of the trained models were evaluated by calculating the precision, recall, F1-score, and ROC-AUC values in the test dataset. All of the procedures were run on Ubuntu 18.04 (DeepStation DK1000, 16 GB RAM, GPU=1).

### Construction of CRE arrays in the tomato genome

The constructed DL model was applied to the 1-kb promoter sequences from the transcription start sites of all the genes in the tomato genome (*N* = 34,066, ITAG4.0; ftp://ftp.solgenomics.net/tomato_genome/annotation/ITAG4.0_release/). We extracted sequence tiles from the promoter region with a sliding window (31-bp bin, and 2-bp step) and input them to the prediction model. The confidence for the prediction was binalized with the threshold = 0.8, then summarized in a 10–50-bp bin to make a one-dimensional binary CRE array per gene for each TF. The resultant CRE arrays on 2,000 randomly selected genes in the tomato genome (Supplementary Table S4) were clustered (or regularized) using a K-means^++^ clustering algorithm (kmeanspp in R) with *K* = 1–150. On the basis of the transition of the sum of squared errors of prediction (Supplementary Fig. S3), we adopted *K* = 50 as the putatively optimized cluster number. The CRE arrays (named by binding TF) with the highest Pearson correlation coefficient to the center array of each cluster, were used for the following expression pattern predictions.

### Deep learning models for predicting expression patterns from CRE arrays

Total 34,066 CREs arrays were annotated with the binary categories in the gene expression pattern in the two criteria (BRup and BRdown). We made a four-fold cross-validation set from all the genes in the tomato genome, allocating 25% for testing and 75% for training/validating samples. For the training/validating samples, we randomly selected 70% for training and 30% for validating. These training/validating sets were applied to one-dimensional convolutional neural network (CNN) models (https://github.com/Takeshiddd/Genome-wide-cis-decoding-for-expression-designing-in-tomato-with-cistrome-and-explainable-deep-lear), which were constructed with the sequential API model of Keras 2.2.4 (https://keras.io/). We examined kernel size (3-20), layer depth (3-16 conv. layers), epoch numbers (5–200), learning rates (0.01–0.000001), optimizers (NAdam, Adam, RMSProp, and SGD), and decay, to optimize the performance in each classification task for at least 40 times. The optimized epoch numbers were defined as the point where additional 10 epochs made no significant reductions in the validation loss. The class weight option (“class_weight” in Keras) was set with the bias in the sample numbers in the two classes. The performances of the trained models were evaluated with the ROC-AUC values in the testing samples.

### Multiple regression and LAMP analyses

Quantitative (or TF-biding site numbers) or binary (or presence/absence of TF-biding sites, with various thresholds) CREs arrays were annotated with the binary categories in the gene expression pattern. Multiple regression test was performed with generalized linear model in R. Limitless-Arity Multiple-testing Procedure (LAMP)^14^ analysis, which is a code for listing up significant combinations of the TFs without an arity limit, was performed to according to the developers instruction (http://a-terada.github.io/lamp/), with Fisher’s exact test.

### Feature visualization in deep learning predictions

An implementation of the feature visualization methods using the iNNvestigate library^47^ has been deposited at https://github.com/uchidalab/softmaxgradient-lrp and https://github.com/Takeshiddd/Genome-wide-cis-decoding-for-expression-designing-in-tomato-with-cistrome-and-explainable-deep-lear. Guided Grad-CAM^17^ and LRP and its variants^48-49^ were implemented to find the important nucleotide residues in CREs. Guided backpropagation^50^ was implemented to reveal the CRE bins relevant to the prediction of expression patterns.

### Vector constructions

To construct the reporter vectors for the transient reporter assay, the intact 1-kb promoter region of the aminocyclopropane-1-carboxylic acid synthase 2 gene (*ACS2*) in the tomato genome (Solyc01g095080) was amplified by PCR from genomic DNA of cv. Micro-Tom using PrimeSTAR GXL DNA Polymerase (TaKaRa) and two primer sets, SlACS2-prom1k-pPLV-F/R or SlACS2-prom1k-TOPO-F/R (Supplementary Table S11). The point-mutated *ACS2mut* allele was artificially synthesized by Eurofins Genomics (Tokyo, Japan), then amplified with the same primer sets. The amplicons from SlACS2-prom1k-pPLV-F/R were cloned into pPLV4^51^ using an In-Fusion HD Cloning kit (Clontech), to construct p*ACS2*-GFPx3 and p*ACS2mut*-GFPx3, where the triplicated green fluorescent proteins (GFPs) were under the control of the *ACS2* or *ACS2mut* promoters, respectively (Supplementary Table S11). The amplicons from SlACS2-prom1k-TOPO-F/R were cloned into the pENTR/D-TOPO cloning vector (Thermo Fisher Scientific) and then cloned into pGWB35^52^ using Gateway LR clonase II (Thermo Fisher Scientific), to construct p*ACS2*-Luc and p*ACS2mut*-Luc, where the firefly luciferase was under the control of the *ACS2* or *ACS2mut* promoters, respectively.

To construct the effector and reference vectors, total RNA was extracted from a ripening fruit pericarp in cv. Eco Sweet, with PureLink Plant RNA Reagent (Thermo Fisher Scientific). The CDS of the *NON-RIPENING* gene (*NOR*, Solyc10g006880) was amplified from the synthesized cDNA by PCR using PrimeSTAR GXL DNA Polymerase (TaKaRa) and the primer set SlNOR-pPLV26-F/R (Supplementary Table S11). The *Renilla* luciferase CDS was amplified from a pRL-null vector (Promega) by PCR using PrimeSTAR GXL DNA Polymerase (TaKaRa) and the primer set RenLuc-pPLV26-F/R (Supplementary Table S11). The amplicons were cloned into pPLV26^50^ using an In-Fusion HD Cloning kit (Clontech) to design p35S-*NOR* and p35S-RenLuc, where the *NOR* and *Renilla* luciferase CDSs were under the control of CaMV35S promoters.

### Transient reporter assay

To assess the activation ability of the *ACS2* and *ACS2mut* promoters in ripening tomato pericarp, we conducted transient dual-luciferase assays with p*ACS2*-Luc, p*ACS2mut*-Luc, pMock-Luc and p35S-RenLuc, which were introduced into *A. tumefaciens* strain EHA105 using the helper vector pSOUP. The transformed agrobacterium was cultured at 28°C for 32 hours, then suspended in Murashige and Skoog (MS) medium (pH 5.3) that contained 20 μg/mL acetosyringone. The concentration was adjusted to OD600 = 2.0. *Agrobacterium* suspension for the negative control (pMock-Luc + p35S-RenLuc), the positive case (p*ACS2*-Luc + p35S-RenLuc), and the mutated case (p*ACS2mut*-Luc + p35S-RenLuc), were inoculated directly into tomato (cv. Eco Sweet) fruit pericarp at the MG, BR and LR stages (6, 12, and 5 biological replicates, respectively) with a 1-ml syringe. Two days after infection, a 10 mm × 10 mm piece of tissue surrounding the inoculation point was applied to the Dual-Luciferase Reporter Assay System (Promega), to detect the firefly luciferase activity (or activation of the *ACS2* and *ACS2mut* promoters), with standardization of the over-expressed *Renilla* luciferase activity. The luciferase luminescence was detected using a ChemiDoc Imaging system (BioRad) and analyzed using Image Lab (BioRad).

To assess the activation ability of the *ACS2* and *ACS2mut* promoters by *NOR*, we conducted transient reporter assays in *Nicotiana benthamiana* leaves with GFP or luciferase as the reporters. For the assay with the GFP reporter, p35S-*NOR*, p*ACS2*-GFPx3, and p*ACS2mut*-GFPx3 were introduced into *Agrobacterium tumefaciens* strain EHA105, as described, and then transiently introduced to the fourth and fifth leaves of *N. benthamiana* plants carrying 8–10 leaves by agrobacterium infiltration. *Agrobacterium* suspension for the control with no effector (p35S-Mock + p*ACS2*-GFPx3), the positive case (p35S-*NOR* + p*ACS2*-GFPx3), or the mutated case (p35S-*NOR* + p*ACS2mut*-GFPx3) were inoculated into the same leaves, with 16 biological replicates. Their relative GFP activities on microscope images were compared under fixed exposure (383 ms) with excitation by the filtered 470– 495 nm laser line. For the dual luciferase assay, p35S-*NOR*, p*ACS2*-Luc, p*ACS2mut*-Luc, and p35S-RenLuc were introduced into *A. tumefaciens* strain EHA105. The transient transformation was conducted as described, with 18 biological replicates. The *N. benthamiana* leaves were harvested 2-days after infection and then applied to a Dual-Luciferase Reporter Assay System (Promega) to detect the activation of the *ACS2* and *ACS2mut* promoters by *NOR*, with standardization of the over-expressed *Renilla* luciferase activity.

## DATA AVAILABILITY

The data that support the findings of this study are available from the corresponding author upon request. All analytical codes and scripts developed in this study have been deposited on GitHub and are publicly available at https://github.com/Takeshiddd/Genome-wide-cis-decoding-for-expression-designing-in-tomato-with-cistrome-and-explainable-deep-lear.

## SUPPLEMENTARY INFORMATION

Supplemental Figures (S1-S9)

Supplemental Tables (S1-S11)

## ACKNOWLEDGEMENTS

We thank Margaret Biswas, PhD, from Edanz Group (https://en-author-services.edanz.com/ac) for editing a draft of this manuscript. This work was supported by PRESTO from Japan Science and Technology Agency (JST) [JPMJPR20Q1] to T.Ak., and Grant-in-Aid for JSPS Fellows for [19J23361] to K.M, JSPS KAKENHI [18H02199] to T.Ak., and [JP16H06280] to S.U.

## AUTHOR CONTRIBUTION

T.Ak., and S.U. conceived the study. T.Ak., T.K., and T.Ar. designed the experiments. T.Ak., K.M., and E.K. conducted the experiments. T.Ak., K.M., E.K., and K.T. analyzed the data, T.Ak., Y.K., and K.U. constructed and maintained the facilities. T.Ak., K.T., and S.U. developed the programs and analytic codes. T.Ak., K.M., E.K., T.K., T.Ar., and S.U. drafted the manuscript. All authors approved the manuscript.

## REFERENCES

1. Charoensawan, V., Wilson, D. & Teichmann, S. A. Genomic repertoires of DNA-binding transcription factors across the tree of life. Nucleic Acids Res. 38, 7364–7377 (2010).

2. Tomato Genome Consortium. The tomato genome sequence provides insights into fleshy fruit evolution. Nature 485, 635–641 (2012).

3. Roulin, A. et al. The fate of duplicated genes in a polyploid plant genome. Plant J. 73, 143–153 (2013).

4. Lynch, M. & Conery, J. S. The evolutionary fate and consequences of duplicate genes. Science 290, 1151–1155 (2000).

5. Wray, G. A. et al. The evolution of transcriptional regulation in eukaryotes. Mol. Biol. Evol. 20, 1377–1419 (2003).

6. Carroll, S. B. Evo-devo and an expanding evolutionary synthesis: a genetic theory of morphological. Cell 134, 25–36 (2008).

7. Kobayashi, S., Goto-Yamamoto, N. & Hirochika, H. Retrotransposon-induced mutations in grape skin color. Science 304, 982–982 (2004).

8. Naito, K. et al. Unexpected consequences of a sudden and massive transposon amplification on rice gene expression. Nature 461, 1130–1134 (2009).

9. Alonge, M. et al. Major impacts of widespread structural variation on gene expression and crop improvement in tomato. Cell 182, 145–161 (2020).

10. Rodríguez-Leal, D., Lemmon, Z. H., Man, J., Bartlett, M. E. & Lippman, Z. B. Engineering quantitative trait variation for crop improvement by genome editing. Cell 171, 470–480 (2017).

11. Li, Q., Sapkota, M. & van der Knaap, E. Perspectives of CRISPR/Cas-mediated cis-engineering in horticulture: unlocking the neglected potential for crop improvement. Hort. Res. 7, 1–11 (2020).

12. Weirauch, M. T. et al. Determination and inference of eukaryotic transcription factor sequence specificity. Cell 158, 1431–1443 (2014).

13. O’Malley, R. C. et al. Cistrome and epicistrome features shape the regulatory DNA landscape. Cell 165, 1280–1292 (2016).

14. Terada, A., Okada-Hatakeyama, M., Tsuda, K. & Sese, J. Statistical significance of combinatorial regulations. Proc. Natl. Acad. Sci. U. S. A. 110, 12996–13001 (2013).

15. LeCun, Y., Bengio, Y. & Hinton, G. Deep learning. Nature 521, 436–444 (2015).

16. Bach, S. et al. On pixel-wise explanations for non-linear classifier decisions by layer-wise relevance propagation. PloS one 10, e0130140 (2015).

17. Selvaraju, R. R. et al. Grad-cam: Visual explanations from deep networks via gradient-based localization. ICCV 618–626 (2017).

18. Zhou, J. et al. Deep learning sequence-based ab initio prediction of variant effects on expression and disease risk. Nat. Genet. 50, 1171–1179 (2018).

19. Akagi, T. et al. Explainable Deep Learning Reproduces a ‘Professional Eye’ on the Diagnosis of Internal Disorders in Persimmon Fruit. Plant Cell Physiol. 61, 1967–1973 (2020).

20. Mejía-Guerra, M. K. & Buckler, E. S. A k-mer grammar analysis to uncover maize regulatory architecture. BMC Plant Biol. 19, 1–17 (2019).

21. Washburn, J. D. et al. Evolutionarily informed deep learning methods for predicting relative transcript abundance from DNA sequence. Proc. Natl. Acad. Sci. U.S.A. 116, 5542–5549 (2019).

22. Wang, H., Cimen, E., Singh, N. & Buckler, E. Deep learning for plant genomics and crop improvement. Curr. Opin. Plant Biol. 54, 34–41 (2020).

23. Tian, Q. et al. MRCNN: a deep learning model for regression of genome-wide DNA methylation. BMC Genom. 20, 1–10 (2019).

24. Alipanahi, B., Delong, A., Weirauch, M. T. & Frey, B. J. Predicting the sequence specificities of DNA-and RNA-binding proteins by deep learning. Nat. Biotechnol. 33, 831–838 (2015).

25. Chow, C. N. et al. PlantPAN3. 0: a new and updated resource for reconstructing transcriptional regulatory networks from ChIP-seq experiments in plants. Nucleic Acids Res. 47, D1155–D1163 (2019).

26. Higo, K., Ugawa, Y., Iwamoto, M. & Korenaga, T. Plant cis-acting regulatory DNA elements (PLACE) database: 1999. Nucleic Acids Res. 27, 297–300 (1999).

27. Smith, C. J. S. et al. Antisense RNA inhibition of polygalacturonase gene expression in transgenic tomatoes. Nature 334, 724–726 (1988).

28. Sheehy, R. E., Kramer, M. & Hiatt, W. R. Reduction of polygalacturonase activity in tomato fruit by antisense RNA. Proc. Natl. Acad. Sci. U.S.A. 85, 8805–8809 (1988).

29. Vrebalov, J. et al. MADS-box gene necessary for fruit ripening at the tomato ripening-inhibitor (Rin) locus. Science 296, 343–346 (2002).

30. Uluisik, S. et al. Genetic improvement of tomato by targeted control of fruit softening. Nat. Biotechnol. 34, 950–952 (2016).

31. Zou, J. et al. A primer on deep learning in genomics. Nature Genet. 51, 12–18 (2019).

32. Bailey, T. L., Williams, N., Misleh, C. & Li, W. W. MEME: discovering and analyzing DNA and protein sequence motifs. Nucleic Acids Res. 34, 369–373 (2006).

33. Shinozaki, Y. et al. High-resolution spatiotemporal transcriptome mapping of tomato fruit development and ripening. Nature Comm. 9, 1–13 (2018).

34. Manning, K. et al. A naturally occurring epigenetic mutation in a gene encoding an SBP-box transcription factor inhibits tomato fruit ripening. Nature Genet. 38, 948–952 (2006).

35. Zhong, S. et al. Single-base resolution methylomes of tomato fruit development reveal epigenome modifications associated with ripening. Nature Biotech. 31, 154–159 (2013).

36. Li, Z. et al. Histone demethylase SlJMJ6 promotes fruit ripening by removing H3K27 methylation of ripening-related genes in tomato. New Phytol. 227, 1138–1156 (2020).

37. Giovannoni, J. J. Genetic regulation of fruit development and ripening. Plant Cell 16, S170–S180 (2004).

38. Weng, L. et al. The zinc finger transcription factor SlZFP2 negatively regulates abscisic acid biosynthesis and fruit ripening in tomato. Plant Physiol. 167, 931–949 (2015).

39. Chung, M. Y. et al. A tomato (Solanum lycopersicum) APETALA2/ERF gene, SlAP2a, is a negative regulator of fruit ripening. Plant J. 64, 936–947 (2010).

40. Liu, M. et al. The chimeric repressor version of an Ethylene Response Factor (ERF) family member, Sl-ERF. B3, shows contrasting effects on tomato fruit ripening. New Phytol. 203, 206–218 (2014).

41. Liu, M. et al. Comprehensive profiling of ethylene response factor expression identifies ripening-associated ERF genes and their link to key regulators of fruit ripening in tomato. Plant Physiol. 170, 1732–1744 (2016).

42. Gao, Ying. et al. Re-evaluation of the nor mutation and the role of the NAC-NOR transcription factor in tomato fruit ripening. J Exp Bot. 71, 3560–3574 (2020).

43. Gao, Y. et al. A NAC transcription factor, NOR-like1, is a new positive regulator of tomato fruit ripening. Hort. Res. 5, 1–18 (2018).

44. Doudna, J. A., & Charpentier, E. The new frontier of genome engineering with CRISPR-Cas9. Science, 346 1258096 (2014).

45. Espley, R. V. et al. Multiple repeats of a promoter segment causes transcription factor autoregulation in red apples. Plant Cell 21, 168–183 (2009).

46. Jakoby, M. et al. bZIP transcription factors in Arabidopsis. Trends Plant Sci. 7, 106–111 (2002).

47. Alber, M. et al. iNNvestigate neural networks!. J. Mach. Learn. Res. 20 1–8 (2019).

48. Montavon, G., Binder, A., Lapuschkin, S., Samek, W. & Müller, K. R. Layer-wise relevance propagation: an overview. in Explainable AI, ser. Lecture Notes in Computer Science. Springer, 11700, 193–209 (2019).

49. Iwana, B. K., Kuroki, R. & Uchida, S. Explaining convolutional neural networks using softmax gradient layer-wise relevance propagation. ICCVW, 4176–4185 (2019).

50. Springenberg, J. T. Unsupervised and semi-supervised learning with categorical generative adversarial networks. arXiv preprint 1511.06390 (2015).

51. De Rybel, B. et al. A versatile set of ligation-independent cloning vectors for functional studies in plants. Plant Physiol. 156, 1292–1299 (2011).

52. Nakagawa, T. et al. Development of series of gateway binary vectors, pGWBs, for realizing efficient construction of fusion genes for plant transformation. J. Biosci. Bioeng. 104, 34–41 (2007).

