## Supplemental data for "Genome-wide *cis*-decoding for expression designing in tomato using cistrome data and explainable deep learning"

**Index:**

Supplementary Figures: S1-S9

Supplementary Tables: S1-S11

**Figure S1**

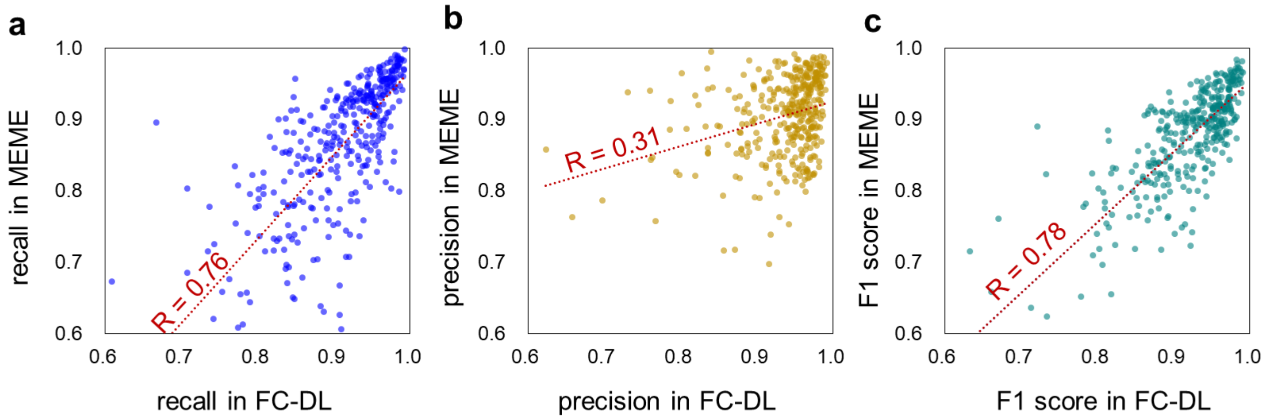


**Figure S1 Correlations for the prediction performances with MEME and the fully-connected deep learning (FC-DL) model.**

MEME and the FC-DL model showed quite high correlations for the performances in the prediction of the CREs from the 370-TFs cistrome datasets. **a.** recall, **b.** precision, and **c.** F1 score. This suggested that intrinsic characteristics of the cistrome datasets, rather than the prediction methods, would have a substantial effect on the quality of the CREs prediction.

**Figure S2**

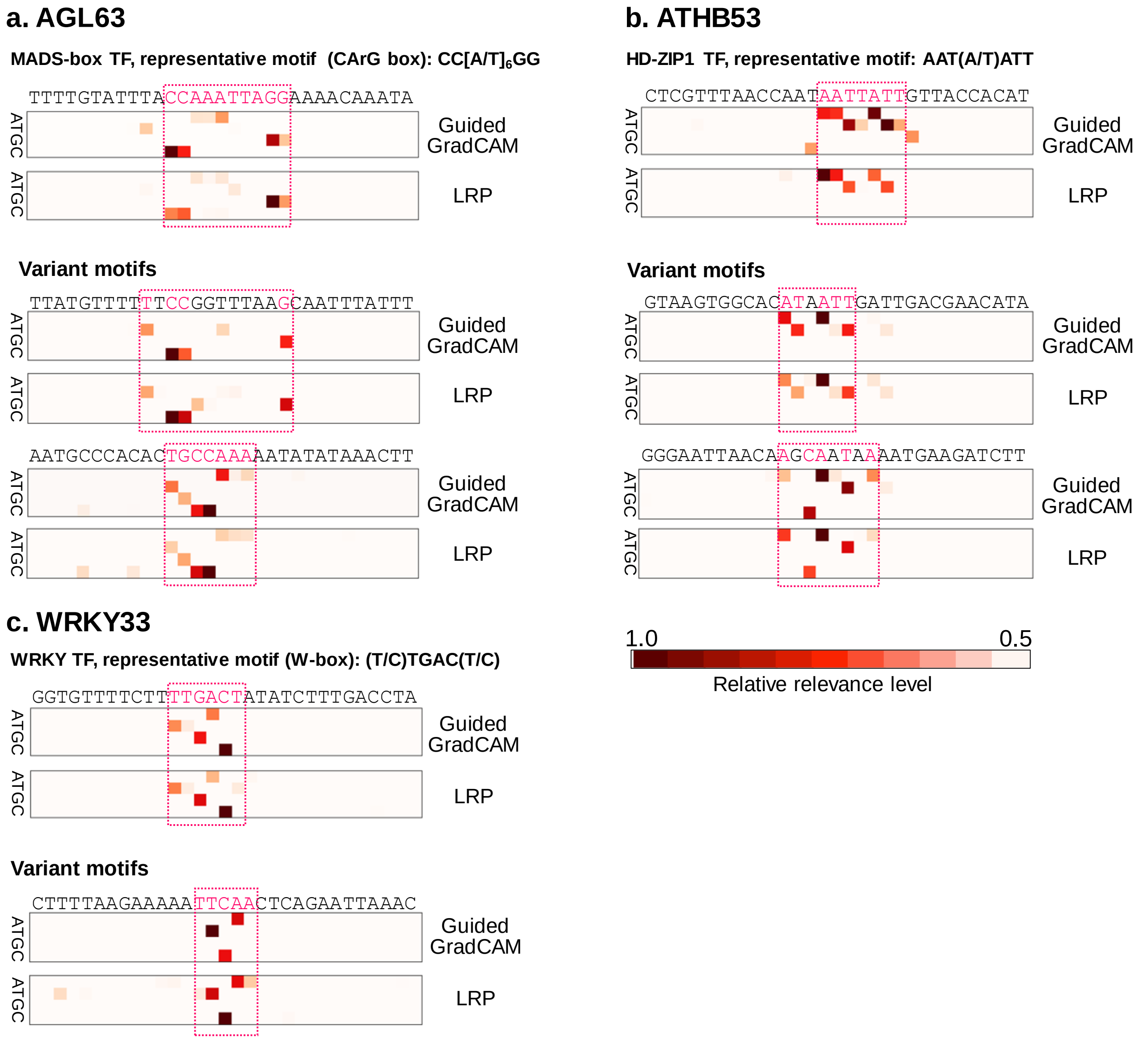


**Figure S2 Feature visualizations in the FC-DL models for three transcription factors.** Detection of the nucleotide residues relevant to the prediction of CREs by the FC-DL model in three TFs, MADS-box AGL63 (**a**), HD-ZIP1 ATHB53 (**b**), and WRKY33 (**c**), with two feature visualization methods. For each TF, TF-binding sequence tiles with high-confidence (>0.95) for the prediction, were exemplified. The relevance was properly focused on the residues consistent with the physiologically validated canonical motifs, the CArG motif for AGL63 (Erdmann et al. 2010), the AAT(A/T)ATT short motif for ATHB53 (Akagi et al. 2020), and the W-box motif for WRKY33 (Zheng et al. 2006). The same feature visualization methods also found some variant (or not registered) motifs including minor gaps or substitutions against the canonical motifs.

**Figure S3**

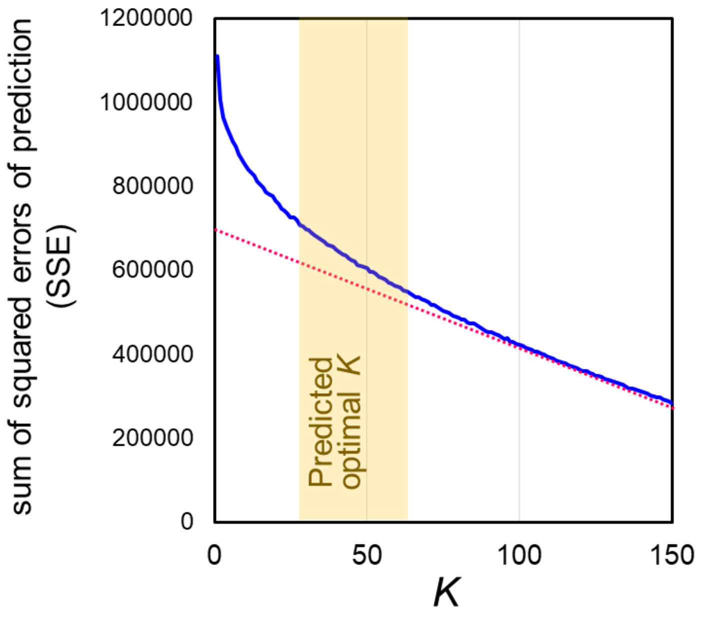


**Figure S3 Estimation of the optimal cluster numbers in Kmeans++**

The sum of squared errors (SSE) of prediction was calculated in K=1~150, with 5 replicates. From the transition of the averaged values of SSE, based on the elbow method, we hypothetically determined K=30~60 for the optimal cluster numbers. In this study, considering the constituting TFs in each cluster, we adopted K=50, as described in the main text.

**Figure S4**

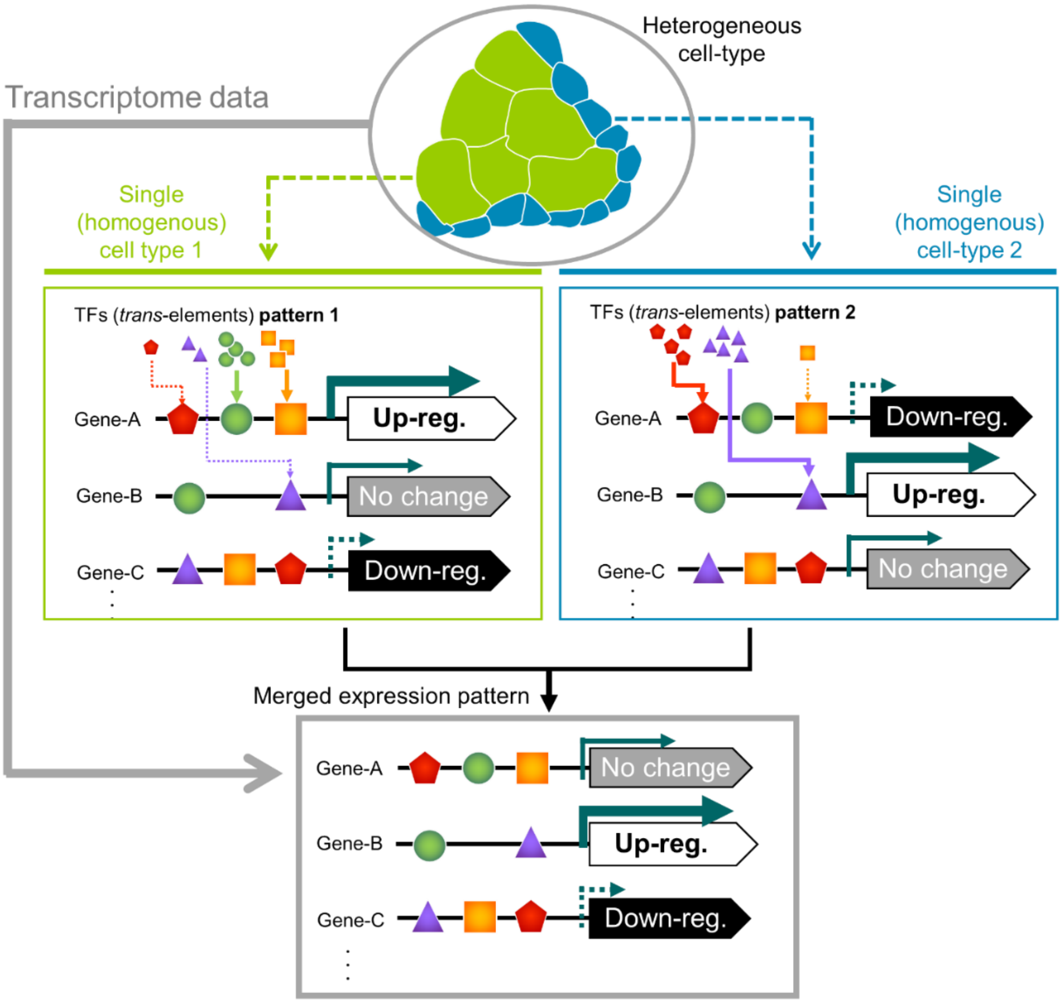


**Figure S4 Difficulty in the prediction of the expression patterns from the CREs, in heterogeneous cell types**

Transcriptome data from heterogeneous cell types would be a result of mixed expression patterns, where the *trans*-elements behaviors are not fixed.

**Figure S5**

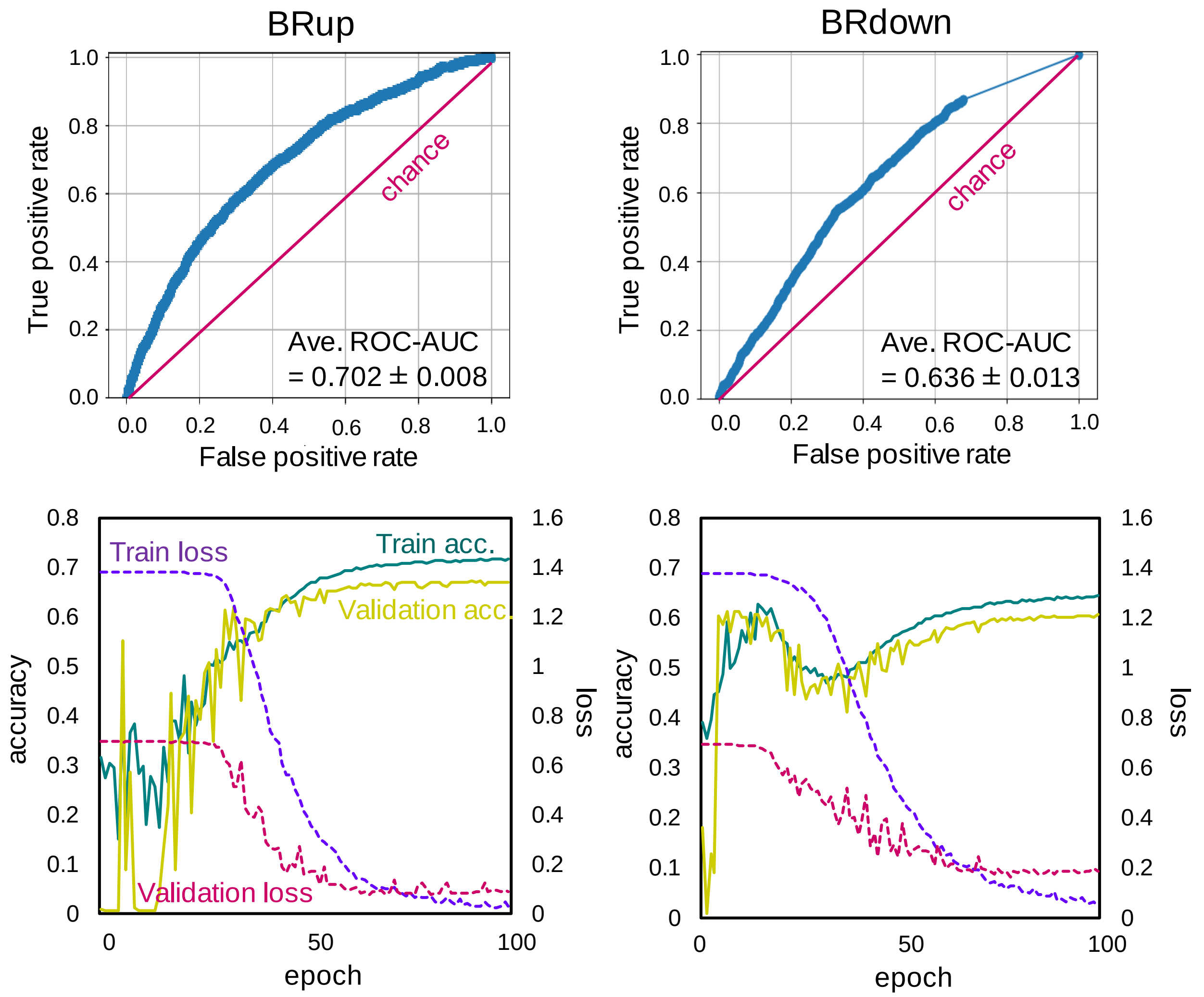


**Figure S5 Learning curves and ROC curves for the BRup and BRdown classifications.**

**a-b.** Examples of the ROC curves for the BRup (**a**) and BRdown (**b**) predictions, in the optimized conditions. **c-d.** Learning curves for the BRup (**c**) and BRdown (**d**) classifications. Transitions of the accuracy (left axis, solid lines) and loss (right axis, dotted lines) were given for the training and validation samples.

**Figure S6**

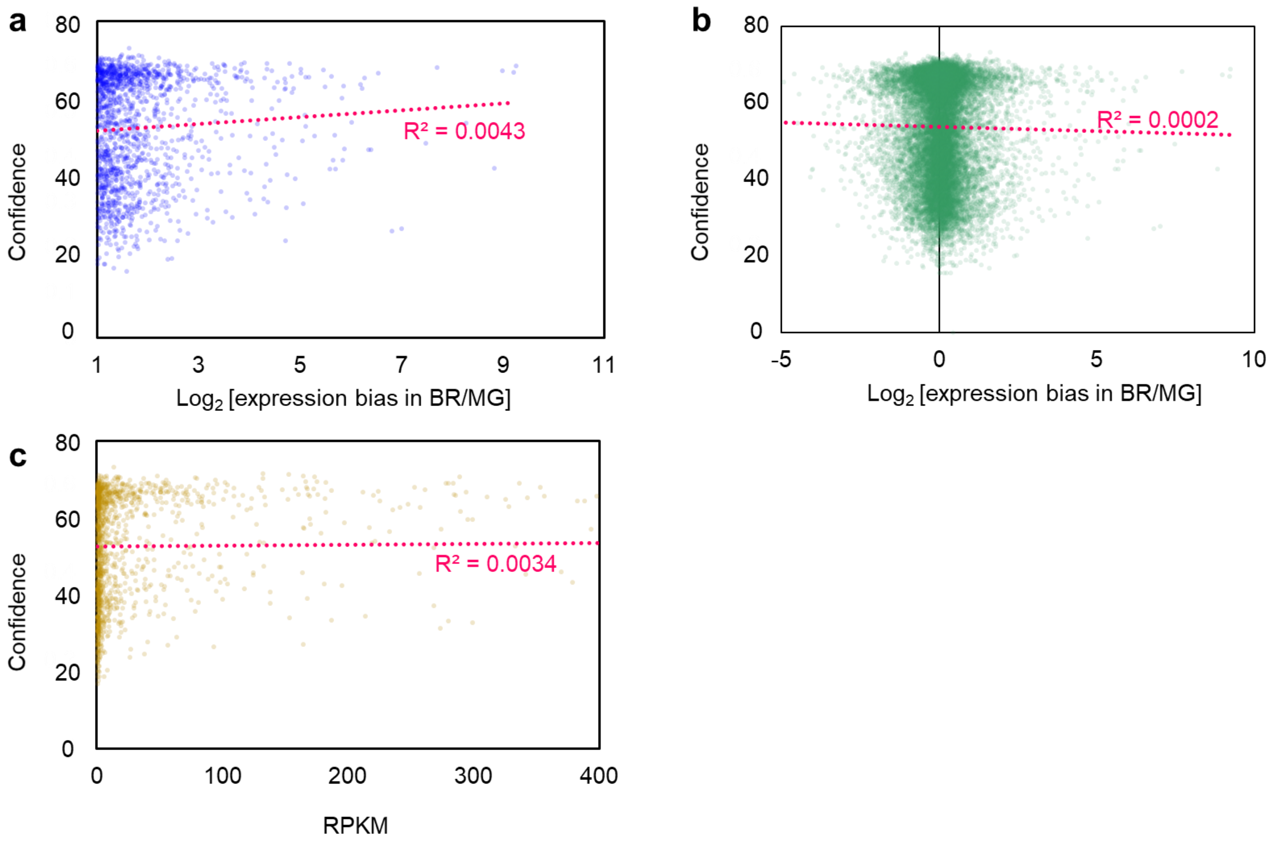


**Figure S6 Correlation in the confidence for the BRup prediction and the expression bias or abundances**

The prediction confidence and the expression bias between BR and MG ($log_2$ values), exhibited no significant correlations both in the BRup gene category (**a**) and in the whole genes (**b**) ($R^2$ = 0.0043 and 0.0002, respectively). Expression abundance (or RPKM, >1.0) in the BRup gene category also exhibited no significant correlation to the prediction confidence (**c**).

**Figure S7**

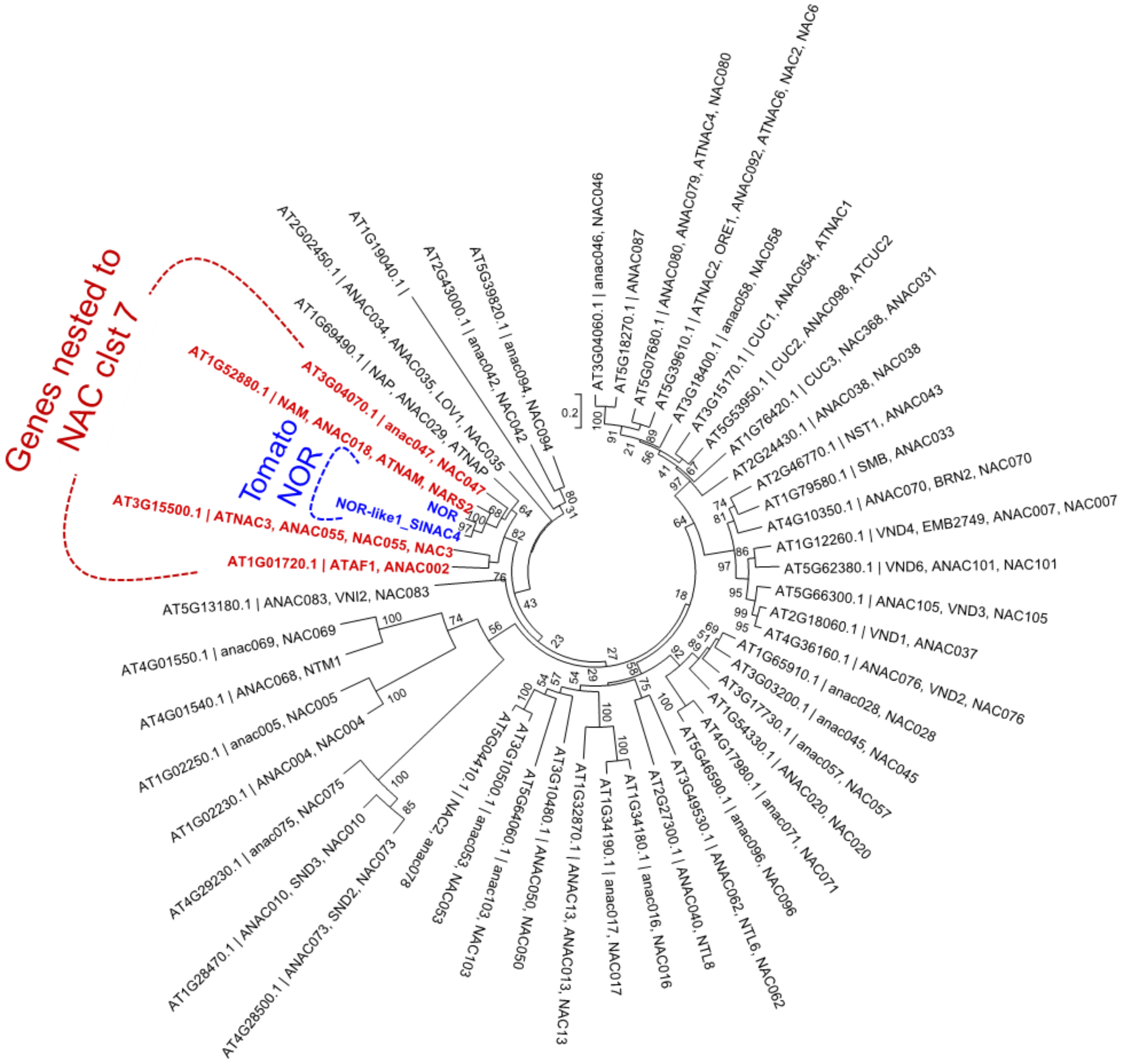


**Figure S7 Phylogenetic relationship of the tomato *NOR* and the Arabidopsis NAC families.** The phylogenetic tree was constructed with amino acid alignments of the tomato NOR/NOR-like 1 (Gao et al. 2018) and the Arabidopsis NAC families used in the cistrome datasets (O'Malley et al. 2016) by the Neighbor-Joining (NJ) method in MEGA X. Tomato NOR/NOR-like 1 genes (highlighted in blue) were nested into the NAC Clst 7 (highlighted in red), with significant support of the bootstrap values (82/100 for the root).

### Figure S8

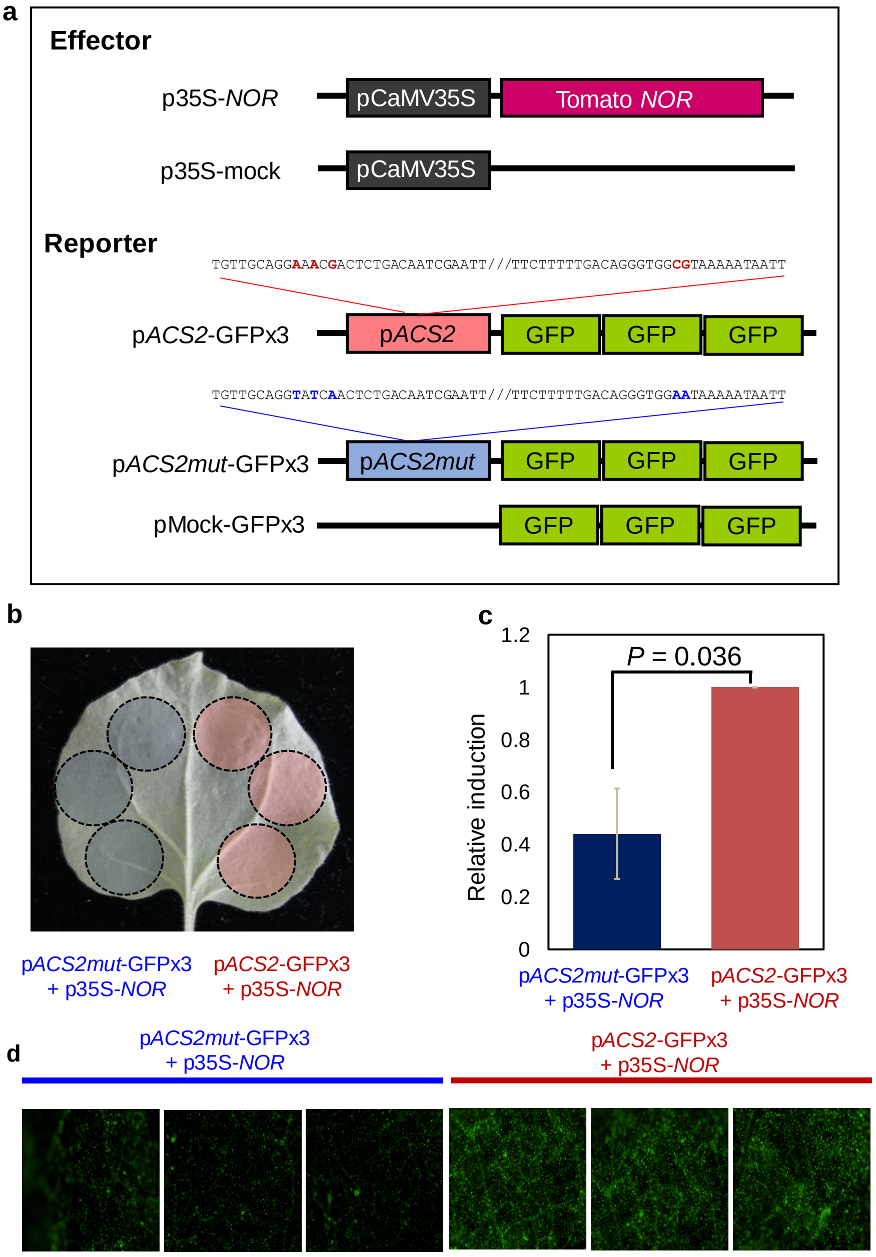


**Figure S8 Visualization of the *ACS2* and *ACS2mut* promoters activation by *NOR*, in *N. benthamiana* leaves.**

**a.** To test in a uniformed condition between *ACS2* and *ACS2mut*, a *N. benthamiana* leaf was infiltrated with *Agrobacterium* including p*ACS2*-GFPx3 and p*ACS2mut*-GFPx3 vectors under the constitutive expression of *NOR*. **b.** The GFP activities were visualized on a microscope under the fixed magnification and exposure (383ms), with excitation by the filtered 470-495nm laser line. Here 6 of the 16 biological replicates were randomly selected. **c.** From the microscope images, the GFP activities were quantified with ImageJ software. The *ACS2mut* promoter showed significantly less GFP induction than the intact *ACS2* promoter (*P* = 0.036, ca. 2.5-fold).

**Figure S9**

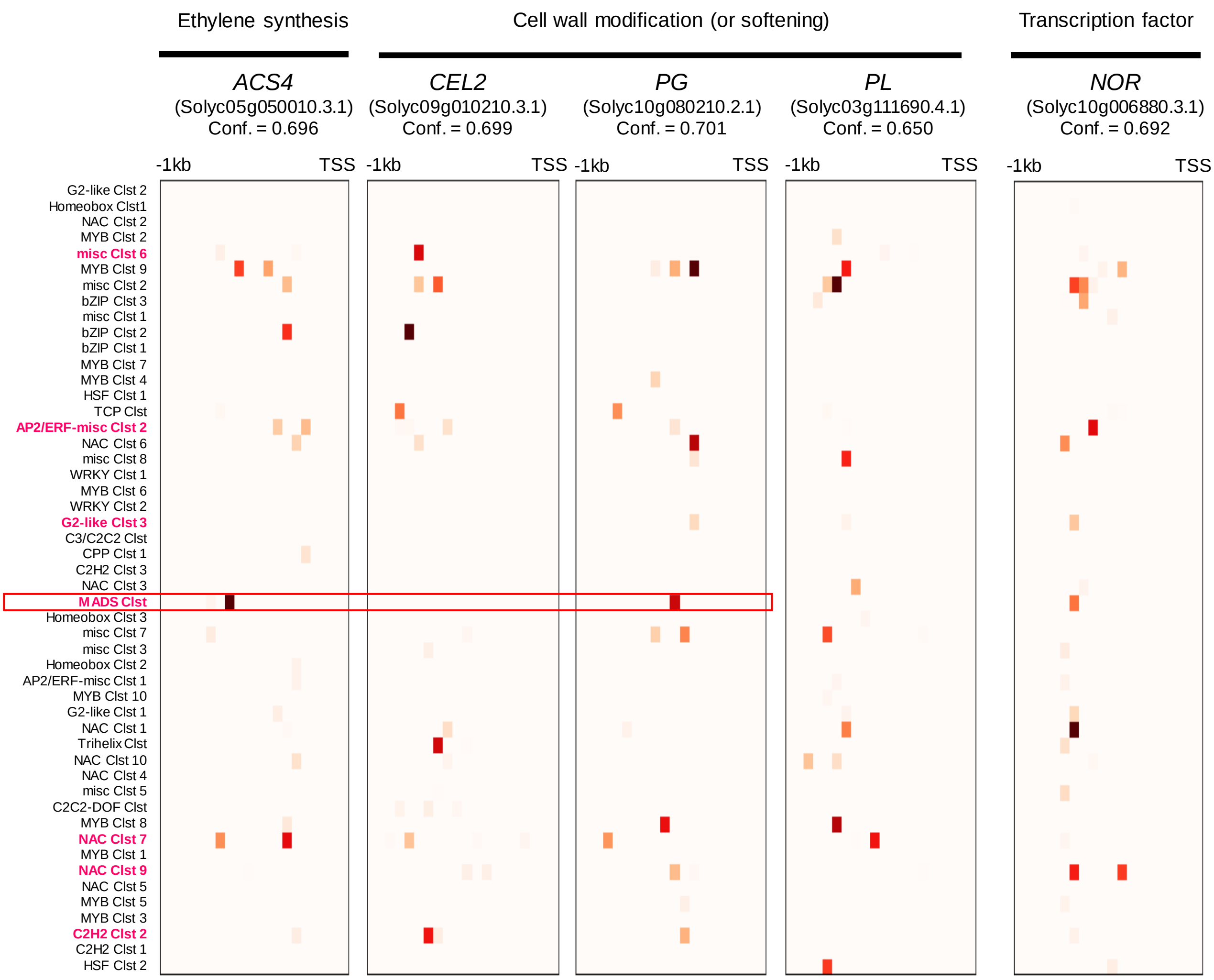


**Figure S9 Identification of the CREs responsible for the BRup in the key genes for tomato fruit ripening.**

The five genes with high-confidence for the BRup prediction (Conf. > 0.65, see Fig. 3c for the confidence distribution) were represented. In the 50 TF clusters, 7 clusters with relatively higher relevance levels (see Fig. 3e-f) were highlighted in pink (on the left side of the panels). As well as *ACS2* (Fig. 3g), *ACS4* and *PG*, which are putatively direct targets of the *RIN* MADS-box TF (Ito et al. 2008, Fujisawa et al. 2012, 2013), exhibited high relevance in the MADS Clst channel, as enclosed in solid red frame. Although conventional techniques, such as yeast one-hybrid (Y1H) method, can find *trans*-factors (or TFs) bound to the promoters of the objective genes, the application of this explainable DL may provide new insights into *trans*-factors directly affecting the targeted expression patterns.

**Supplementary Table S1-S11: See the appendix excel sheet.**

**Table S1** Annotations of the cistrome (DAP-seq) data used in this study.

**Table S2** ROC-AUC values for the CREs classifications in 370 TFs.

**Table S3** Classification abilities in the FC-DL and MEME, among the 370 TFs.

**Table S4** Gene list of which CREs patterns in the promoter regions were applied for the Kmeans++ clustering.

**Table S5** Fifty CREs clusters defined by Kmeans++

**Table S6** Gene list of the BRup category

**Table S7** Gene list of the BRdown category

**Table S8** Multiple regression analysis with the clustered CREs patterns, for the BRup binary classification. The clustered CREs with high relevance in the DL model (see Fig. 3e) were highlighted in bold.

**Table S9** LAMP analysis with the clustered CREs patterns, for the BRup binary classification. The clustered CREs with high relevance in the DL model (see Fig. 3e) were highlighted in bold.

**Table S10** Confidences for the BRup prediction in the representative DEGs related to ethylene production/signaling or ripening in tomato. The genes with the highest 10% confidences for the BRup were highlighted in bold. *ACS2* was highlighted in red.

**Table S11** Primer note for this study.

**REFERENCES**


Akagi, T. et al. The persimmon genome reveals clues to the evolution of a lineage-specific sex determination system in plants. *PLoS Genet.* **16**, e1008566 (2020).

Erdmann, Robert, et al. "GORDITA (AGL63) is a young paralog of the Arabidopsis thaliana Bsister MADS box gene ABS (TT16) that has undergone neofunctionalization." *Plant J.* 63, 914-924 (2010).

Fujisawa, M. et al. Direct targets of the tomato-ripening regulator RIN identified by transcriptome and chromatin immunoprecipitation analyses. *Planta* **235**, 1107-1122 (2012).

Fujisawa, M. et al. A large-scale identification of direct targets of the tomato MADS box transcription factor RIPENING INHIBITOR reveals the regulation of fruit ripening. *Plant Cell.* **25**, 371-386 (2013).

Gao, Y. et al. A NAC transcription factor, NOR-like1, is a new positive regulator of tomato fruit ripening. *Hortic Research* **5**, 1-18 (2018).

Ito, Y. et al. DNA - binding specificity, transcriptional activation potential, and the rin mutation effect for the tomato fruit - ripening regulator RIN. *Plant J.* **55**, 212-223 (2008).

O'Malley, R. C. et al. Cistrome and epicistrome features shape the regulatory DNA landscape. *Cell* **165**, 1280-1292 (2016).

Zheng, Z. et al. Arabidopsis WRKY33 transcription factor is required for resistance to necrotrophic fungal pathogens. *Plant J.* **48**, 592-605 (2006).
